# Hybrid Gaussian Process Models for continuous time series in bolus fed-batch cultures

**DOI:** 10.1101/2021.12.27.474269

**Authors:** M. Nicolás Cruz-Bournazou, Harini Narayanan, Alessandro Fagnani, Alessandro Butté

**Affiliations:** DataHow AG, Zürichstrasse 137, 8600 Dübendorf, Switzerland; Technische Universität Berlin, Bioprocess Engineering, Ackerstrasse 76, 13355 Berlin, Germany; DataHow AG, Zürichstrasse 137, 8600 Dübendorf, Switzerland

**Keywords:** Nonparametric methods, Nonlinear system identification, Grey box modelling, Time series modelling, Gaussian Process Models, Bioprocess Engineering, Mammalian Cell Cultures

## Abstract

Hybrid modeling, meaning the integration of data-driven and knowledge-based methods, is quickly gaining popularity among many research fields, including bioprocess engineering and development. Recently, the data-driven part of hybrid methods have been largely extended with machine learning algorithms (e.g., artificial neural network, support vector regression), while the mechanistic part is typically using differential equations to describe the dynamics of the process based on its current state. In this work we present an alternative hybrid model formulation that merges the advantages of Gaussian Process State Space Models and the numerical approximation of differential equation systems through full discretization.

The use of Gaussian Process Models to describe complex bioprocesses in batch, fed-batch, has been reported in several applications. Nevertheless, handling the dynamics of the states of the system, known to have a continuous time-dependent evolution governed by implicit dynamics, has proven to be a major challenge. Discretization of the process on the sampling steps is a source of several complications, as are: 1) not being able to handle multi-rate date sets, 2) the step-size of the derivative approximation is defined by the sampling frequency, and 3) a high sensitivity to sampling and addition errors. We present a coupling of polynomial regression with Gaussian Process Models as representation of the right-hand side of the ordinary differential equation system and demonstrate the advantages in a typical fed-batch cultivation for monoclonal antibody production.

## 1. INTRODUCTION

Mathematical representations of (bio)chemical reaction kinetics in complex process systems can be broadly classified into two approaches: (i) data-driven, statistical, black box, or Machine Learning (ML) models, and (ii) first principles-based, mechanistic, or white box models Narayanan et al. (2020b). Hybrid models or grey-box models have emerged aiming to combine the features of existing modeling techniques and very interesting applications have been reported in chemical engineering and in biotechnology Zhang et al. (2019); Von Stosch et al. (2014).

### 1.1 Hybrid/Grey-Box Models

A typical procedure to setup the hybrid model involves formulating the basic mass and energy balances while using data-driven models to describe complex unknown properties of the system. Some commonly used data-driven models are: artificial neural network Tian et al. (2001), Adaptive Regression Splines Duarte and Saraiva (2003), Support Vector Machine (for regression) Hu et al. (2011), and genetic programming Marenbach and Bettenhausen (1997).

### 1.2 Gaussian Process State Space Models

A very promising class are Gaussian Process State Spaces Models (GPSSM) Hewing et al. (2018), which aim to predict the dynamics of a process and its time evolution by describing the state space with Gaussian Process Models (GPM) Rasmussen (2003). By this, the advantages of GPMs can be applied to nonlinear dynamic processes with a very limited knowledge of the first principles governing the time evolution of the system such as fed-batch cultivation Hutter et al. (2021). The added value of GPSSMs has been documented in several applications Kocijan (2016), and offer a clear advantage on systems, with small to mid size data sets with noise that is close to Gaussian, and where the uncertainty in the predictions play an important role. Still, a major drawback is the forward problem handling of the data and its representation, and issues associated with it (e.g. sensitivity to noise). An additional limitation is the impossibility to treat multi-rate data sets properly. Finally, the discretization of the continuous dynamic process is bound to the sampling rate of the data set, which can pose several difficulties in scarce sampling processes (e.g. mammalian cell cultures with samplings on a daily basis).

### 1.3 Continuous Gaussian Process State Space Models

We present an alternative approach in an effort to enable training of the GPSSMs considering a continuous time system and exploiting the advantages of the inverse problem theory. To achieve this, we use a polynomial regression on the states over time and train the GPMs on its derivatives. In a sense, the rationale is close to formulating a parameter estimation problem using full discretization Cervantes and Biegler (2009) if we consider that the right hand side of the differential equation system is defined by GPMs. As a result, we obtain the exact derivatives over time for any desired point and are able to train significantly more efficient and flexible GPSSMs. One of the major challenges is the fact that polynomial slopes are not known. To tackle this, we apply a slight modification of the IPDA method as presented by Varziri et al. (2008). We demonstrate that the increase in computational burden due to the iterative procedure and larger number of support points, as well as the complexity of the problem formulation due to additional hyperparameters and optimization to be solved, is justified for many cases.

### 1.4 Hybrid Modelling of Bioprocesses

The inherent complexity of cellular biochemical reaction that take place in bioprocess cultivations combined with the limited and low-quality observations obtained from laboratory experiments and production sites, has proven to be an ideal playground for the use of hybrid models. On the one hand, thousands of biochemical reactions are taking place in each one of the millions of cells inside the reactor. On the other, it is very challenging to observe most of the species (e.g. intracellular metabolites, enzymes, ribosomes) involved in the process. Even the desired product and its quality require typically complex and sophisticated analytical methods, strongly limiting the number of possible samples per experiment (mostly end point evaluations). On the one hand, mechanistic models require model development and specific experiments to support it. On the other, Black box models require many experiments as the relation between inputs and outputs that is typically very non-linear. Both approaches are infeasible due to intrinsic costs of each experiment, such that hybrid approaches are essential to tackle this situation. Without loss of generality, we demonstrate the advantages of the continuous

GPSSMs in a cell culture fed-batch process for production of a Monoclonal Antibody (MAB) Li et al. (2010). These processes are known to be very difficult to model with first principle formulations and require large adaptations even for slight changes in the process, cell line, or the product. For many of the industrial process parameters that are used to improve the cultivation, (e.g. pH and temperature) there exist no mechanistic equations. Finally, the cost of each experiment is very high, such that only a few experiments are possible limiting to the information that can be obtained about the system.

## 2. METHODOLOGY

### 2.1 Gaussian Process State Space Models

For the sake of generality, we stick to the definition of GPSSM given in Eleftheriadis et al. (2017) The GPSSM is here considered as a dynamical system whose building blocks are Gaussian processes. The dynamical system is represented by

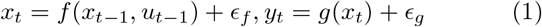

where *t* indexes time, *x* ∈ *R*^*D*^ is a latent state, *u* ∈ *R*^*P*^ are control signals (actions) and *y* ∈ *R*^*O*^ are observations. Furthermore, we assume i.i.d. Gaussian system/measurement noise. The state-space model equation (1) can be fully described by the measurement and transition functions, *g* and *f*, respectively. The key idea of a GPSSM is to model the transition function *f* in equation (1) using GPMs:

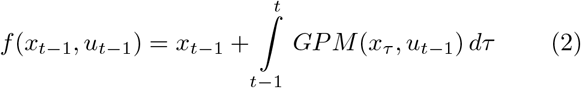

### 2.2 cGPSSM

The applications of GPSSM are well documented, the reader is referred to Kocijan (2016) for a detailed description and case studies. Still, especially for dynamic systems with a low frequency sampling rate, the limitations of GPSSMs become clear. The step size of the prediction Δ(*x*_*t*_−*x*_*t−*1_) is fixed and defined by the sampling intervals. For this reason the discretization is inaccurate if the sampling is not faster than the time constant of the dynamic system. Furthermore, the highest sampling rate that can be used for the training of the GPs is subject to the fixed rate where all observed states are measured. These are important limitations that give Differential Equation Systems a significant advantage for a number of applications.

We propose an alternative approach based on three iterative steps: 1. fitting polynomial curves to the data set, 2. training the GPs with the derivative of the polynomials, and 3. penalizing the polynomial fit if there is a large difference between the GP predicted derivatives and the polynomial derivatives

As a result, GPs are trained with the exact derivative at any given point and can be used for adaptive multistep integration methods. Furthermore, the penalization assures that the derivatives of all polynomials fitted in the different data sets comply with the function defined by the GP. This allows, considering that the data set is sufficiently large, a well-posed problem penalizing divergent dynamics in different data sets.

This approach stems from the gradient matching method, were a similar methodology is suggested for parameter inference of differential equation systems using GPs instead of solving the inverse problem. *“Techniques based on gradient matching, which aim to minimize the discrepancy between the slope of a data interpolant and the derivatives predicted from the differential equations, offer a computationally appealing shortcut to the inference problem”* Dondelinger et al. (2013).

## 3. ALGORITHM

### 3.1 Polynomial Regression on Finite Elements

In order to cope with the nature of bioprocesses, piece wise polynomial functions divided in finite elements are preferred in terms of simplicity and flexibility. Furthermore, in many applications, mammalian cell cultures are performed using a bolus feeding strategy causing discontinuities in the state variable profiles due to the sudden change in concentration at each pulse. The grid should be set such that the value of all states is known in the initial time point of each finite element. This is typically the same at the rate of one finite element per day in mammalian cell cultures. For the sake of simplicity, we do not consider the case where some values are missing at the start of the finite elements. This issue can easily be tackled by selecting other methods to initialize the polynomials. Here, a polynomial regression is used to generate the initial derivative values to be used in the training of the GPM for each finite elements. There are of course several polynomial types that can be used, spline functions, or Lagrange polynomials for example. In this first case we use a simple quadratic equation of the form:

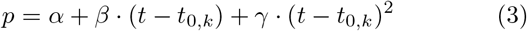

where *α, β, γ* are the polynomial coefficients, *t* represents the time and *t*_0,*k*_ the initial time point of the *k* − *th* FE. In the first iteration, *γ* is set to zero, resulting in a linear step identical to the GPSSM formulation.

We determine the coefficients of the quadratic polynomials, minimizing the weighted least squares function:

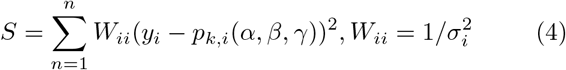

### 3.2 GPM training with polynomial derivatives

Once the polynomials have been fitted to the data, we can use their derivatives as a first approximation to train the GPMs:

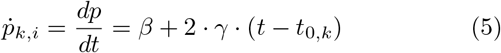

The GPMs are trained using the state variables, the experimental set points, and the time varying control variables (typically piece wise constant) as inputs. The outputs 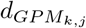 are the derivatives of the polynomials on the selected support points 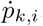.

It is worth stressing out that the polynomials in the first iteration are restricted to *γ* = 0, hence the GPMs are firstly trained with a piece wise constant profile. Therefore, selecting a proper Kernel is crucial to ensure a convergence towards the actual curvature of the time evolution of the states. Considering that we are dealing with processes properly described by Lipschitz continuous functions, the GPMs approximate the derivatives of the polynomials closer to the average value of all experiments. As a result, the derivatives are dragged towards the “real” continuous curvature of the process profile. The linearization enforced on known curves (caused by the large step discretization) delivers during training different derivative values for very similar input variables. The GPMs predict, hence the mean slope averaging the inconsistent slopes throughout the state space.

### 3.3 Iterative convergence to continuous functions

In order to drive the polynomials to the expected curve, the iterative solution of the least square problem in equation (4) is penalized by the mean derivative predicted by the GPMs at the given input conditions:

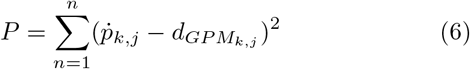

The resulting function for optimization being:

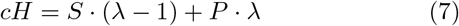

As shown in figure 1 through an iterative process, the polynomials converge to a proper description of the true dynamics of the states at any point inside the design space, given following conditions:

- a sufficiently informative data set,
- a correct selection of the finite element size,
- the proper degree and type of polynomials, and
- an observable system with the sampled state variables

**Fig. 1.**
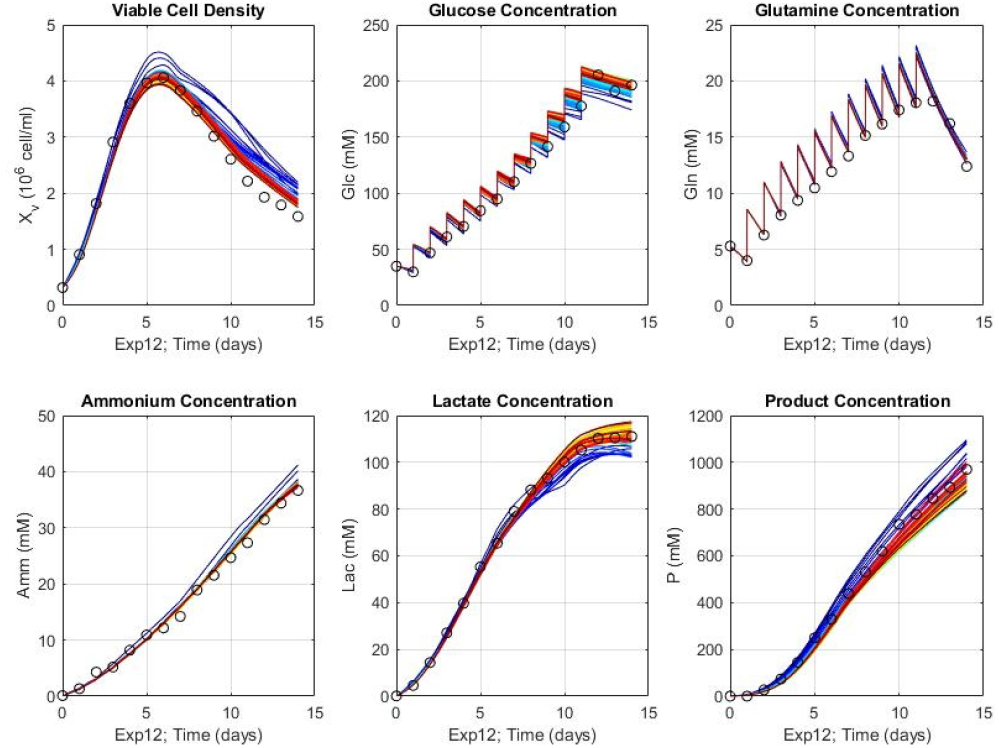
Simulation results for an experiment of the training subset. The simulation of all 60 iterations of the cGPSSM framework. The first iteration is represented by the dark blue line and red represents the last one. The convergence to a significantly better prediction and a smooth representation of the time evolution of the states is clearly represented

### 3.4 Numerical integration

Being able to predict the exact derivatives of the function at any given point within the design space, the GPM predictions can now be used to predict the time evolution of the process through numerical approximation. In this example, we use an explicit Euler method with a fixed step size of 0.5 hours. This is sufficient given the smoothness of the process within each finite element and avoids an extensive number of function calls. It is here noteworthy that the pulses are solved with the equality constraints between each finite elements. Depending of the size of the data set contained in the GPM, the prediction calls can rapidly increase the computational burden of the integration. This is especially the case, when standard integrators are used which tend to start with very small step sizes and perform a number of evaluations to assure robustness in the solution. Note that the prediction results now deliver values at a much higher frequency than the sampling one (1 sampling per day), and could also work with adaptive step approaches if necessary. The use of polynomials to link GPMs and data allows a continuous and smooth prediction of the systems dynamics throughout the state space and accordingly of the evolution of the process over time.

Furthermore, due the property of GPMs, the predicted derivatives also account for its uncertainty. This can be used to compute not only the predicted time evolution but also its associated uncertainty. This of course requires a special treatment of the uncertainty propagation during integration, which is out of the scope of this contribution.

### 3.5 Materials

The computations for the selected case study were performed in a simple laptop computer. Memory: 7,6 GiB, Processor: Intel® Core™ i7-8565U CPU @ 1.80GHz × 8, OS: Linux, Ubuntu 18.04.6 LTS. The code was written and computed using Matlab2021a. The packages (toolboxes) used were: the nonlinear least-squares solver lsqnonlin from the optimization toolbox, the polyfit function from the curve fitting toolbox, and the rgp class from the statistical toolbox. The kernel used for the GPMs was the ardexponential kernel with a (*d*+1)−*by*−1 vector *ϕ*, where *ϕ*(*i*) contains the length scale for predictor *i*, and *ϕ*(*d* + 1) contains the signal standard deviation. *d* is the number of predictor variables. Default initial value of the length scale parameters are the standard deviations of the predictors and the signal standard deviation is the standard deviation of the responses divided by square root of 2. That is,

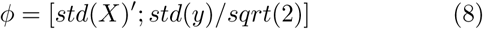

The code can unfortunately not be shared due to Intellectual Property concerns. However, pseudocode is presented in the following section to illustrate the step-by-step procedure for implementation of the proposed algorithm.

#### Algorithm 1

pseudocode for the cGPPM approach: training

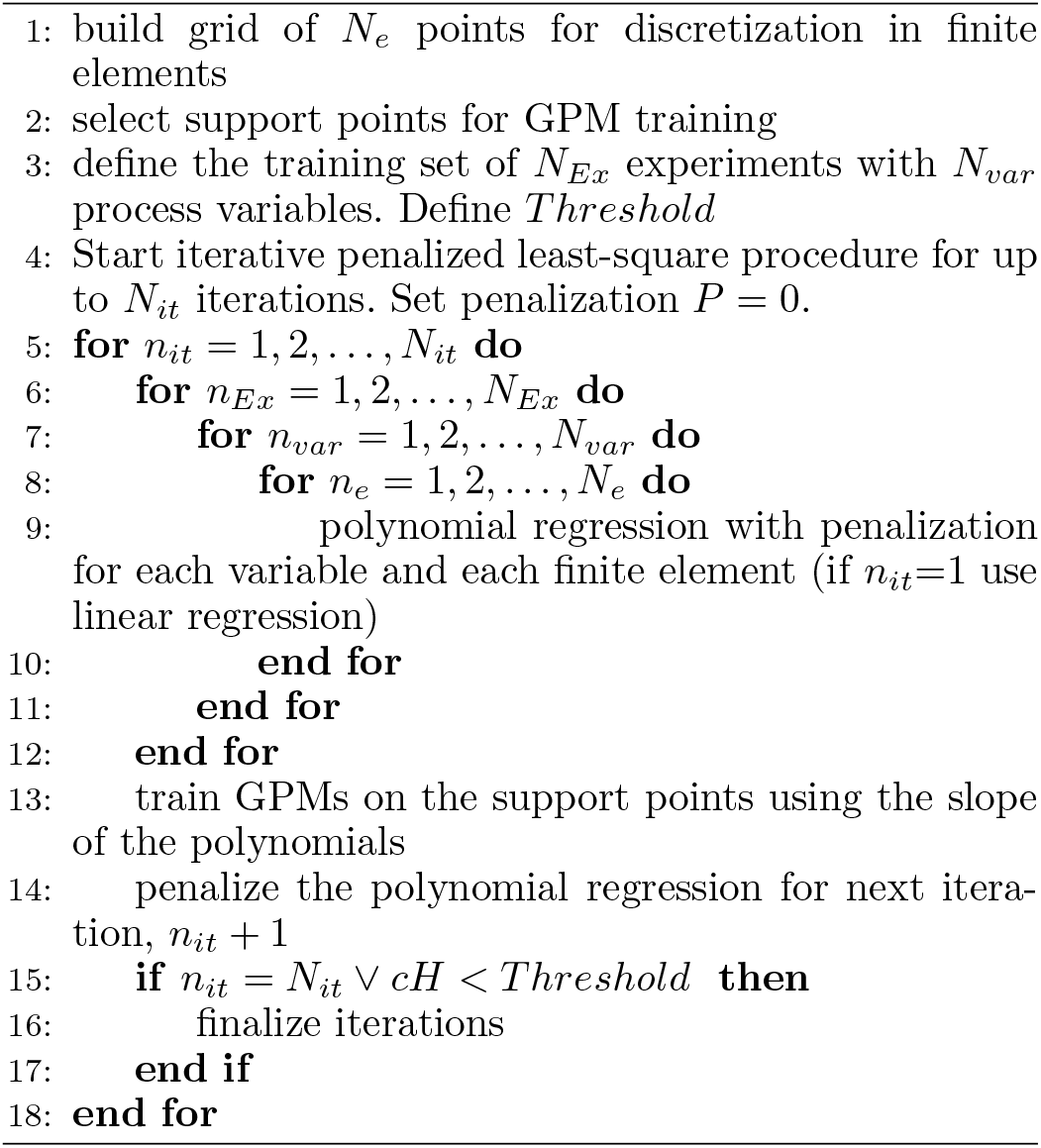

#### Algorithm 2

pseudocode for the cGPPM approach: simulation

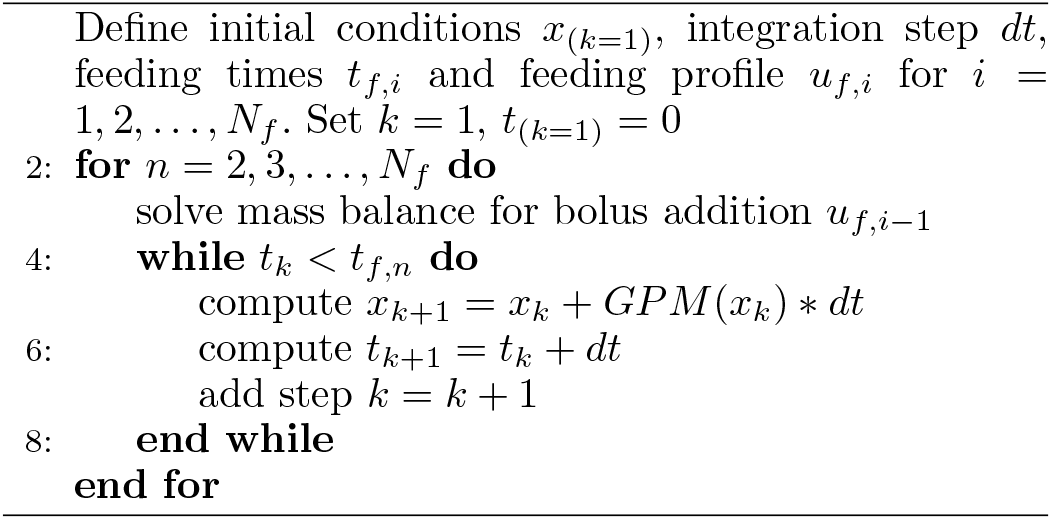

## 4. RESULTS

### 4.1 Case Study

We demonstrate the performance of the approach with a mammalian cell culture bioreactor using a macro-kinetic model to generate in-silico data. A cell culture fed-batch bioreactor is simulated using a system of ODEs with six variables, namely the Viable Cell Density (VCD), glucose as the main substrate (Glc), glutamine (Gln), ammonia (Amm), lactate (Lac) and the product titer. 30 experiments are planned using a Latin hypercube sampling method, varying the bolus feed (F) and the initial concentration of biomass and substrate. 20 experiments are used to train the model and 10 experiments are used to test the model. Time profiles are simulated in the interval of [0, 14] days and measurements perturbed with 15% Gaussian noise. For a detailed description of the insilico data, the reader is referred to Narayanan et al. (2020a). For the evaluation of the results, concentrations normalized to the maximum value of the respective states in the training set are used as inputs. The different models are compared based on the Root Mean Squared Error in prediction (RMSEP), computed with respect to the true values (not the perturbed measurements), of individual states.

### 4.2 Description capabilities of the cGPSSM

As we can see in Figure 1, the resulting model can describe a continuous smooth dynamic profile of the fed-batch process. It can be seen, that the iterative procedure converges to a smooth representation that is indeed closer to the “real” trajectory of the process. The model catches the smooth evolution seen in the Cell Viable Density but also (due to the proper grid discretization) the discrete changes in the glucose profile. The discontinuities caused by the sudden additions are handled in the switch from one finite element to the next one. Figure 2 shows a comparison between the standard method (discretized GPM) versus cGPSSM on the validation data set demonstrating the improved prediction capabilities of the proposed approach. Furthermore, the smooth curves, describing the real time evolution of the process, that are generated with this novel method are depicted in Figure 3. An important concern is the convergence of the penalized LSQ to a stable solution, which was indeed the case in our example (see Figure 4). Still, the method can fail if the value of *λ* is selected to close to 1, or the insufficiency of data impede an accurate prediction for each given state.

**Fig. 2.**
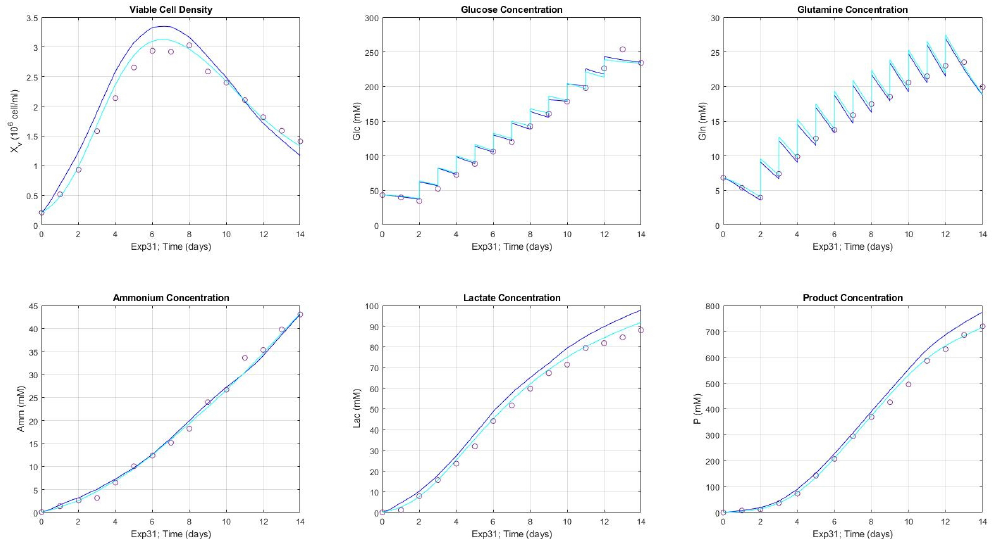
Simulation results on an experiment of the validation subset showing first (standard GPSSM solution) and 15th iterations of the cGPSSM framework. The six state variables are depicted. The better prediction of a previously unseen experimental setup is demonstrated.

**Fig. 3.**
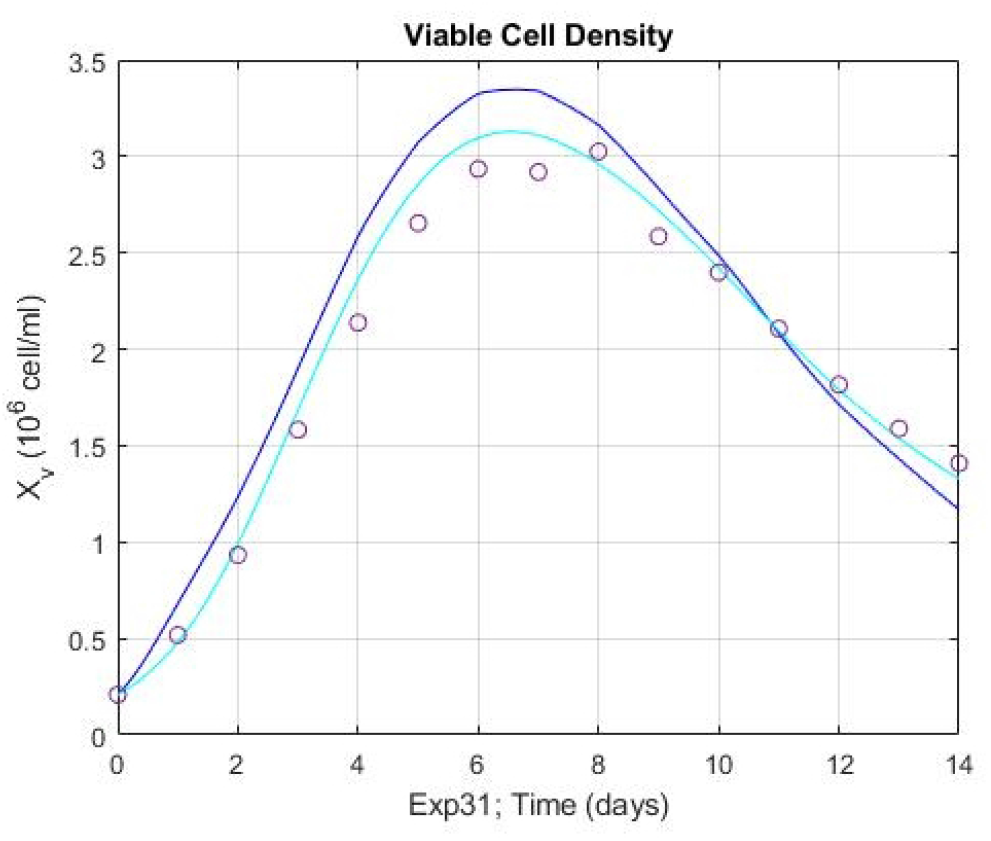
Simulation results of the Viable Cell Density (VCD) time evolution in an experiment of the validation subset showing first (standard GPSSM solution) and 15th iterations of the cGPSSM framework. The better prediction of a previously unseen experimental setup is demonstrated.

**Fig. 4.**
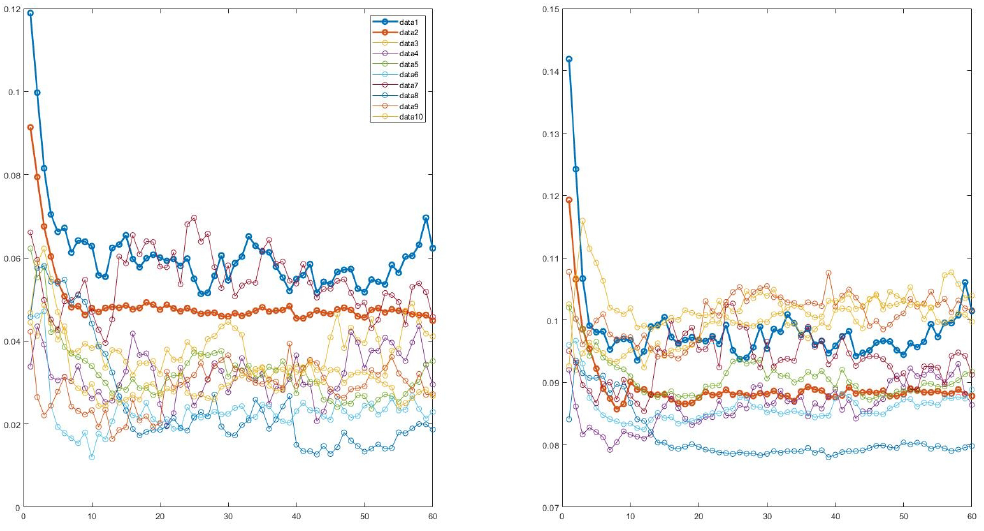
Evolution of the Root Mean Square Error (RMSE) throughout the iterations, for experiments in the training subset (left) and in the validation subset (right). For the specific case, 15 iterations are sufficient to reach the final result and best predictions.

Two important conditions are to be considered, that are only valid in the in-silico data set and not necessarily in real process data. First, the system we are dealing with is time invariant and fully observable. There are, hence, no complications due to state variables that affect the process but are not monitored and the dynamics of the system never change (in contrast to real biological systems). Secondly, the noise is Gaussian by which completely fulfills the underlying assumptions and the necessary conditions to use GPMs. Real data sets do not have a perfectly Gaussian noise, and, in fact, the error distributions can be very complex. Still, several real applications are close to the conditions given in our example and the capability of GPMs to describe mammalian cell cultures has been widely reported Tsopanoglou and del Val (2021).

## 5. DISCUSSION

Despite their advantages, there still some challenges related to the implementation of the approach. Clearly, the iterative process represents a significantly higher computational burden, and it adds further hyperparameters to the framework. Additionally, the integration can also be computational demanding, especially if the uncertainty intervals are computed using ensemble models. Furthermore, there is no experimental evidence on the true curvature of the polynomial between two sampling points. As a result, it is difficult to validate the final results besides visual inspection. Still, this approach allows the use of GPSSMs in multi-rate sampling data sets enabling the implementation of GPM technologies in new fields and applications.

## 6. CONCLUSIONS

In this work, we present an alternative formulation to obtain Gaussian Process State Space Models with a smooth continuous time evolution of the state variables. To achieve this, we use polynomials as interpolants in the data and solve a penalized least-squares problem to enforce consistency across data from different experiments. The mammalian cell culture example demonstrates not only the good performance of the approach, but also its added value for the description of complex processes that are widely used in biotechnology and biopharma. The bolus fed-batch process is described precisely in comparison to the ground truth obtained from the in-silico simulations and a continuous smooth time evolution is obtained despite the large gaps in the measurement samplings. This approach can also overcome important issues that standard machine learning algorithms face. First, it can deal with process information with different sampling rates in its varibales. Second, it allows the use of adaptive step integrators and delivers an accurate prediction at any given time point within the design space. Thirdly the convergence of the iterative training and penalized least-square procedure assures a consistent and somewhat “well-posed” problem.

## Notes

### Competing Interest Statement

The authors have declared no competing interest.

